# Endothelial ADAM17 Promotes Neutrophil Migration and Pulmonary Microvascular Permeability in ARDS

**DOI:** 10.64898/2026.01.21.700786

**Authors:** Anna Biedritzky, Carolin Kleinmaier, Anika Fuhr, Kristian-Christos Ngamsri, Franziska Konrad, Michael Koeppen

**Affiliations:** Department of Anesthesiology and Intensive Care Medicine, University Hospital of Tuebingen, Germany; Center for Anesthesiology, Intensive Care Medicine, Perioperative Medicine and Pain Therapy Markgröningen, RKH regional clinics holding and services GmbH

**Author notes:** Correspondence to: Franziska M. Konrad, MD Klinik für Anästhesiologie Orthopädische Klinik Markgröningen Kurt-Lindemann-Weg 10 71706 Markgröningen OR Michael Koeppen, MD Department of Anesthesiology and Intensive Care Medicine University of Tuebingen Hoppe-Seyler-Strasse 3 72076 Tuebingen Phone: 0049/7071-2986935 Fax: 0049/7071-2986935.

**Keywords:** Acute respiratory distress syndrome, Endothelial ADAM17, Microvascular permeability, Endothelial barrier dysfunction, Neutrophil migration, Pulmonary inflammation

## Abstract

Acute respiratory distress syndrome (ARDS) is characterized by profound endothelial barrier disruption, excessive neutrophil recruitment, and sustained pulmonary inflammation. A Disintegrin and Metalloproteinase 17 (ADAM17) regulates inflammatory signaling through ectodomain shedding of adhesion molecules and cytokine receptors, yet its endothelial-specific contribution to ARDS remains poorly defined.

We identify endothelial ADAM17 as a central regulator of vascular permeability, neutrophil trafficking, and inflammatory amplification in LPS-induced acute pulmonary inflammation. LPS markedly increased pulmonary ADAM17 expression, whereas endothelial-specific ADAM17 deletion reduced total lung ADAM17 mRNA by 77.5%. Endothelial ADAM17 promoted disruption of endothelial junctions and protein-rich pulmonary edema by modulating JAM-A and VE-cadherin. Concomitantly, endothelial ADAM17 facilitated neutrophil transmigration into interstitial and alveolar compartments through altered expression of PSGL-1 and CD49d.

Mechanistically, endothelial ADAM17 enhanced TNF receptor 1 and IL-6 receptor signaling, increasing proinflammatory mediator release. Pharmacological ADAM17 inhibition recapitulated the protective phenotype of endothelial ADAM17 deficiency, attenuating neutrophil recruitment and preserving endothelial barrier integrity.

These findings establish endothelial ADAM17 as a key driver of inflammatory vascular dysfunction in ARDS and support ADAM17 as a rational therapeutic target.

## Introduction

Acute respiratory distress syndrome (ARDS) is a rapidly progressive and often fatal condition characterized by non-cardiogenic pulmonary edema, profound hypoxemia, and widespread alveolar inflammation. Despite advances in supportive care, mortality in severe ARDS remains high, reaching up to 40% (1). At the core of ARDS pathophysiology is a sustained hyperinflammatory response that disrupts the alveolar–capillary barrier, increases microvascular permeability, and drives neutrophil infiltration into the lung parenchyma.

Neutrophils and endothelial cells engage in dynamic interactions during acute inflammation. While neutrophils mediate pathogen clearance, their excessive accumulation contributes to tissue damage and worsens pulmonary edema. However, the molecular mechanisms governing neutrophil recruitment and endothelial barrier dysfunction in ARDS remain incompletely understood.

A Disintegrin and Metalloproteinase 17 (ADAM17) is a membrane-bound protease that modulates inflammation through the ectodomain shedding of adhesion molecules and cytokine receptors, including TNF-α, L-selectin, and junctional adhesion molecule A (JAM-A). Prior studies have implicated ADAM17 in leukocyte trafficking and barrier regulation across various inflammatory settings, and its pharmacological inhibition has shown promise in preclinical models of lung injury. However, ADAM17 has complex, context-dependent roles, exerting both pro- and anti-inflammatory effects depending on cellular origin and timing, which limits the utility of global inhibition.

Importantly, the specific contribution of endothelial ADAM17 to pulmonary inflammation and barrier dysfunction remains poorly defined. Since endothelial cells form the first interface for leukocyte transmigration and actively contribute to inflammatory signaling, dissecting the role of ADAM17 in this compartment is critical to understanding the pathogenesis of ARDS and identifying viable therapeutic targets.

Here, we tested the hypothesis that endothelial ADAM17 promotes neutrophil recruitment, disrupts endothelial barrier integrity, and amplifies pulmonary inflammation in ARDS. Using a conditional, endothelial-specific ADAM17 knockout mouse model and two pharmacological inhibitors (TAPI-1 and GW280264X), we investigated how endothelial ADAM17 modulates microvascular permeability, neutrophil migration, inflammatory signaling, and cytokine release in a murine model of LPS-induced acute lung injury.

## Material and methods

### Animals

The Adam17^tm1.2Bbl^/J floxed mice (strain: 009597; RRID:IMSR_JAX:009597, MGI ID: MGI:4354847) were obtained from Jackson Laboratory (Bar Harbor, USA). This mouse line contains loxP sites flanking exon 2 of the ADAM17 locus. To generate the tissue-specific knockout, these mice were crossed with Tie2-Cre mice (Tg(Tek-cre)12Flv; MGI ID: MGI: 2136412) kindly provided by Prof. Dr. Peter Rosenberger, resulting in the B6.129P2-ADAM17tm1.2Bbl-Tek-cre1Ywa/J also referred to as ADAM17^fl/fl^ Tie2-Cre^+^. The crossing of these lines produced a tissue-specific ADAM17 knockout in murine endothelial cells via Cre-mediated deletion of the floxed ADAM17 gene. The Cre-negative littermates were used as control animals and will be referred to as ADAM17^fl/fl^ Tie2-Cre^-^. Female and male animals were used in a 1:1 ratio at eight to twelve weeks age. All animal protocols were approved by the Animal Care and Use Committee of the University of Tuebingen (A05/21G) and adhered to the principles of the 3Rs (Replacement, Reduction, Refinement) to ensure ethical standards.

### Murine model of LPS-induced acute pulmonary inflammation and drug administration

Acute pulmonary inflammation was induced via LPS inhalation (0,5 mg/ml in 7 ml sterile saline, derived from *Salmonella enterica* serotype enteritidis, Sigma Aldrich) for 30 minutes in a custom-built chamber. Nebulized LPS resulted in a transient pulmonary inflammation characterized by increased neutrophil migration, elevated microvascular permeability and altered chemokine release. For systemic ADAM17 inhibition in ADAM17^fl/fl^ Tie2-Cre^-^, TAPI-1 (5 µg/g body weight) and GW280264X (30 µg/g body weight) were administered intraperitoneally one hour before LPS inhalation.

Identical ADAM17^fl/fl^ Tie2-Cre^-^ control animals were used for comparison to both endothelial ADAM17 deletion and systemic inhibition, in accordance with the 3Rs principle from Russell and Burch (1959) to minimize animal numbers. For transparency, this is noted in the relevant figure legend.

### Immunofluorescence

Paraffin-embedded lung sections were deparaffinized, rehydrated, and subsequently fixed in 4 % PFA. After washing, tissue sections were permeabilized with 1 % Triton X-100 and blocked with 5 % bovine serum albumin (BSA) in PBS to reduce nonspecific binding. Sections were then incubated with primary antibodies targeting ADAM17 (rabbit anti-ADAM17, polyclonal, Bioss, USA), Ly6G/C (rat anti-Ly6G/C, clone RB6-8C5, Abcam, UK), VE-Cadherin (goat anti-VE-Cadherin, polyclonal, R&D Systems, USA), TNFR1 (mouse anti-TNFR1, clone H-5, Santa Cruz, USA), IL-6Rα (mouse anti-IL-6Rα, clone H-7, Santa Cruz, USA), and vWF (sheep anti-vWF, polyclonal, Abcam, UK) followed by incubation with the corresponding fluorescently labelled secondary antibodies including Alexa Fluor 488 goat anti-rabbit, Alexa Fluor 647 goat anti-rat, Alexa Fluor 488 donkey anti-goat, Alexa Fluor 594 goat anti-mouse, Alexa Fluor 488 donkey anti-mouse and Alexa Fluor 647 donkey anti-sheep. Nuclear counterstaining was performed using Roti-Mount FluorCare containing DAPI (4’,6-diamidino-2-phenylindole; HP20.1, Carl Roth, Germany). Images were acquired using a Leica Stellaris 8 confocal microscope and processed with LasX software. For each experimental condition, three independent experiments were performed, each including two technical replicates.

### Flow cytometry-based in vivo migration assay

Neutrophil migration into the inflamed lung structures – vascular, adherent to pulmonary endothelium, interstitial, and bronchoalveolar – was determined by flow cytometry 24 hours after LPS inhalation. PMNs attached to the pulmonary endothelium were tagged by an allophycocyanin (APC)-conjugated anti-Ly6G antibody (clone 1A8, BioLegend, USA), which was injected intravenously into the tail vein immediately prior to anaesthesia. In deep anaesthesia, thoracotomy was performed and followed by blood collection and flushing PBS through the pulmonary vasculature to remove circulating PMNs. Alveolar PMNs were collected via bronchoalveolar lavage. Lung tissue was collected, enzymatically digested and homogenized. Bronchoalveolar lavage (BAL) was centrifuged and cells were resuspended in staining buffer. The whole blood and cell suspensions of lung and BAL were stained with PerCP-conjugated CD45 (clone 30-F11, BioLegend) and PE/Cy7-conjugated Ly6G (clone 1A8, BioLegend) antibodies. Endothelial adherent PMNs were identified as PerCP-CD45^+^/PE/Cy7-Ly6G^+^/APC-Ly6G^+^, interstitial PMNs as PerCP-CD45^+^/PE/Cy7-Ly6G^+^/APC-Ly6G^-^, and bronchoalveolar PMNs as PerCP-CD45^+^/PE-Cy7-Ly6G^+^ (Supplementary Figure 1). Additionally, the surface expression of the adhesion molecules CD162, CD49d and JAM-A as well as ADAM17 was assessed using the following antibodies: PE-conjugated CD162 (clone 2PH1, BioLegend), APC-conjugated CD49d (clone R1-2, BioLegend), PE-conjugated CD321 (clone 27-9, BioLegend), and PE-conjugated ADAM17 (polyclonal, BioLegend).

### Bicinchoninic acid assay for total protein determination

The total protein amount in BAL supernatants was measured as an indirect indicator of microvascular leakage. For this purpose, the BCA assay kit (Pierce Thermo Scientific, USA) was used according to the manufactureŕs instructions.

### Evans blue extravasation assay

For the determination of endothelial barrier dysfunction and microvascular leakage, Evans blue was injected intravenously into the tail vein at a dose of 20 µg/g body weight, five hours and 30 minutes after LPS-inhalation. After 30 minutes incubation, mice were anesthetized, blood collected and subsequently perfused with PBS to remove residual blood from the pulmonary circulation. Lung were excised, weighed, and diluted with PBS. Tissue was homogenized using ceramic beads and homogenates were mixed 1:2 with formamide and incubated at 60 °C for 12-16 hours. Blood samples were centrifuged, plasma was collected and stored at 4 °C overnight. At the following day, lung samples were centrifuged and optical density was determined at 620 nm and 740 nm as reference wavelength. Evans blue concentrations in plasma and lung were determined by regression analysis using the standard curve.

### Immunohistochemistry

To evaluate neutrophil accumulation in the lung 24 hours after LPS exposure, mice were anesthetized and subsequently perfused to remove residual blood from the pulmonary circulation. Lungs were inflated and fixed via tracheal cannulation with 4 % formaldehyde for 10 minutes at a pressure of 25 cm H2O, excised, embedded in paraffin, and sectioned at thickness of 3 µm. For immunohistochemical analysis, sections were first blocked with Avidin solution (Vector Laboratories, USA) and then incubated overnight at 4 °C with a rat anti-mouse Ly6G/C antibody (1:500, clone RB6-8C5, Abcam, UK). After washing, sections were incubated for 1 hour with a biotinylated anti-rabbit IgG secondary antibody (BA400, Vector Laboratories) and visualization with 3,3’-diaminobenzidine (DAB) substrate (SK-4100, Vector Laboratories). Nuclear counterstaining was performed with Mayeŕs hematoxylin (1.09249, Sigma Aldrich, Germany). Images were acquired with a Leitz DM IRB microscope (Leica) and processed using AxioVision software (v4.8.2), while quantitative image analysis was carried out with ImageJ (v1.53k).

### Gene expression

Total RNA was extracted from murine lung tissue three hours following LPS inhalation using TRIzol reagent (Invitrogen, Carlsbad, USA) in accordance with the manufactureŕs protocol. Subsequently, complementary DNA (cDNA) was synthesized from the isolated RNA employing the iScript cDNA synthesis kit (Bio-Rad, Munich, Germany) following the manufactureŕs instructions. Quantitative real-time PCR was then performed to determine relative gene expression levels using the following primers: ADAM17 (5’-gag agc cat cag ttt g-3’, 5’-ccc tcc atg agt ttg c-3’), VE-Cadherin (5’-gtg ctc tcc aca aag ctc gg-3’, 5’-gga gga gct gat ctt gtc cg-3’), iRhom2 (5’-gac acc ttc gac tcc tcc tt-3’, 5’-ccg ctg tat tct ttt agc ggg -3’), AT1R ( 5’-tgc cgt gaa ctt gaa ggt tat tc-3’, 5’-gct tga gaa cac caa gca gc-3’), iTAP (5’-tcc aga act gga ggc ctg a-3’, 5’-ctt ctg ccc cct cca tct tg-3’), TNFR1 (5’-ctt cag cac ccc agg ctt ta-3’, 5’-tcg caa ggt ctg cat tgt ca-3’), TNFR2 (5’-cta agt gtc ctc ctg gcc aat -3’, 5’-tgg gtt ttc aag gcg cag ta-3’), TRAF2 (5’-gct act gct cct tct gcc tg-3’, 5’-gca ggt tct cag tct cca cc-3’), MAPK1 (5’-tga agt tga aca ggc tct ggc -3’, 5’-tcc tct gag ccc ttg tcc tga -3’), MAPK3 (5’-gta cgg cat ggt cag ctc ag-3’, 5’-ctg agg atg tct cgg atg cc-3’), Nf-κB (5’-gcc aag tgc acg agt c-3’, 5’-gaa gga cga gac gag c-3’), p38 (5’-agc tgg tag cgg tgc gat aa-3’, 5’-cat ctc tct ctg tcc agt gcc a-3’), JNK (5’-agc cgt ctc ctt tag cac ag-3’, 5’-tgt atc cga ggc caa agt cg-3’), FADD (5’-ctc aat cgc ctt ccg cat tg-3’, 5’-aca gcc agg tga gaa atg gg-3’), TRADD (5’-tct gca ggt tcg aag ttc cc-3’, 5’-gtg gcc ggt tca cta cga g-3’), CREB (5’-gag cag aca acc agc aga gt-3’, 5’-tgg cat gga tac ctg ggc ta-3’), IL-6R (5’-tca ctg tgc gtt gca aac ag-3’, 5’-gat ccg gct gca cca ttt tt-3’), JAK1 (5’-gcc tgt cta ctc cat gag cc-3’, 5’-gtc tgg atc ttg cct ggt ca-3’), STAT3 (5’-tca gcg aga gca gca aag aa-3’, 5’-ctt ggt ctt cag gta cgg gg-3’), TRIF (5’-ctt ccc cac agt ccc aat cc-3’, 5’-gaa cca tct ggg cat ggt ga-3’), TRAM (5’-tct atg ctt gtc ctc gca cc-3’, 5’-gga gcg act act ttc cca gg-3’).

### ELISA

Levels of the inflammatory mediators TNF-α, CD62L, IL-10, ELA2, and CXCL2/3 were quantified in BAL three hours after LPS inhalation. ELISAs were performed with kits from R&D Systems (USA) according to the manufactureŕs protocol. Levels of soluble VE-Cadherin were evaluated using the kit from Abcam (UK).

### MPO

The release of myeloperoxidase, serving as an indicator of neutrophil activity, was quantified in the supernatant of BAL at 24 hours following LPS inhalation. Quantification of MPO activity was performed using a colorimetric assay, based on the peroxidase-mediated conversion of 2,2’-Azino-di(3-ethylbenzthiazolin-6-sulfonsäure) (ABTS), with optical density determined at 405 nm using a microplate reader (Infinite M200 PRO, Tecan, Switzerland).

### Statistical analysis

Statistical analyses were performed using GraphPad Prism software (v10.1, GraphPad Software, San Diego, USA). Unless stated otherwise, data are presented as mean ± standard deviation (SD). Outlier detection was conducted using the Robust Regression and Outlier Removal (ROUT) method with a coefficient of Q = 1 %. The normality of data distribution was evaluated using both the D’Agostino-Pearson test and the Shapiro-Wilk test. For normally distributed data, group comparisons were carried out using one-way analysis of variance (ANOVA), followed by Bonferronís post hoc test. In cases where data did not meet normality assumptions, the non-parametric Kruskal-Wallis test was applied. All statistical tests were two-tailed, and values of p < 0.05 were considered statistically significant.

## Results

### ADAM17 expression is upregulated during LPS-induced acute pulmonary Inflammation

ADAM17 expression was quantified in murine lung tissue after LPS exposure. Under physiological conditions, vehicle-treated ADAM17^fl/fl^ Tie2-Cre^-^ and ADAM17^fl/fl^ Tie2-Cre^+^ knockout mice showed no differences in pulmonary ADAM17 mRNA expression (Figure 1A). Three hours after LPS inhalation, ADAM17 mRNA levels in the lungs of ADAM17^fl/fl^ Tie2-Cre^+^ knockout mice were reduced by 77.5 % compared to controls, reflecting effective knockout efficiency. At the protein level, statistical analysis of ADAM17 expression in lung tissue (Figure 1B), along with representative z-stack immunofluorescence images (Figure 1C–D), confirmed a significant reduction in ADAM17 protein in knockout mice 24 hours after LPS inhalation.

**Figure 1:**
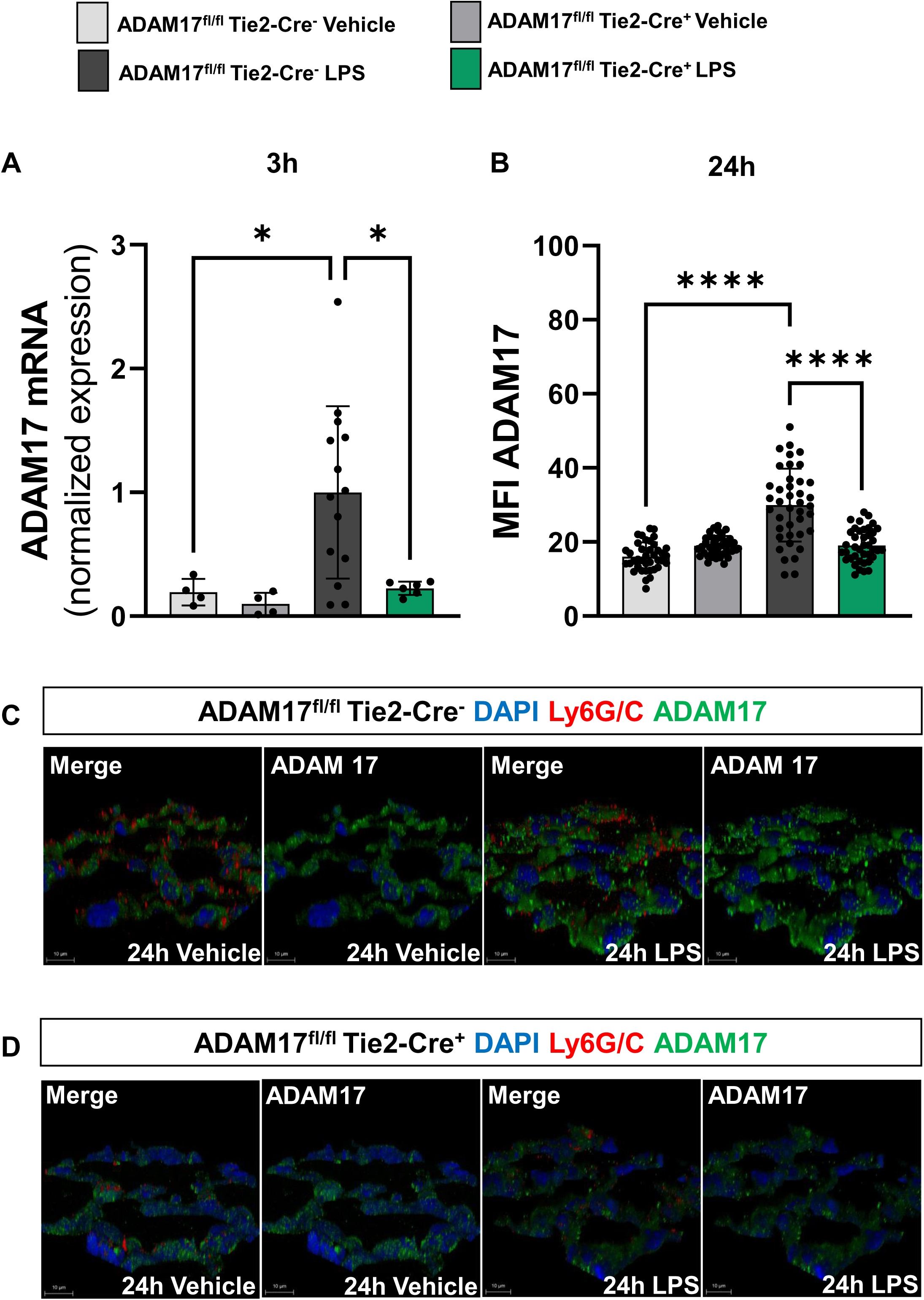
ADAM17 expression is upregulated during LPS-induced acute pulmonary inflammation. **(A)** Gene expression of ADAM17 in lung tissue three hours after LPS inhalation, measured by RT-qPCR (n = 4/4/14/8 mice per group). Expression data were normalized to the ADAM17^fl/fl^ Tie2-Cre^−^ LPS control group, whose mean value was set to 1. **(B)** Quantification of ADAM17 mean fluorescence intensity (MFI) in lung tissue using LasX software (n = 4 mice per group; 10 measurements per image). **(C)** Representative z-stack immunofluorescence images showing ADAM17 expression in lung tissue under the indicated conditions (original magnification ×63). Data are presented as mean ± SD. Statistical analysis was performed using one-way ANOVA with appropriate post hoc testing; *P < 0.05, **P < 0.01, ***P < 0.001, ****P < 0.0001.

These results highlight the pivotal contribution of endothelial-derived ADAM17 to total pulmonary expression. The endothelial-specific deletion was validated by flow cytometry (Supplementary Figure 2). ADAM17 surface expression on pulmonary endothelial cells was quantified by mean fluorescence intensity (MFI) (Supplementary Figure 2A). In ADAM17^fl/fl^ Tie2-Cre^+^ mice, endothelial ADAM17 surface levels were significantly reduced compared to ADAM17^fl/fl^ Tie2-Cre^-^ controls, confirming efficient targeting of endothelial cells. In contrast, ADAM17 surface expression on neutrophils was unchanged across blood, lung, and BAL fluid, and on platelets from blood and lung tissue. However, ADAM17 expression on platelets recovered from BAL fluid was increased, consistent with platelet activation within the inflamed alveolar compartment (Supplementary Figure 2B–C). Importantly, ADAM17 expression on endothelial cells was selectively reduced, confirming the endothelial specificity of the genetic deletion.

### Endothelial ADAM17 promotes vascular barrier disruption during lung injury

To investigate the role of endothelial ADAM17 in vascular permeability, we assessed endothelial junction integrity and lung barrier function following LPS exposure using multiple complementary assays (Figure 2). These included analysis of VE-Cadherin mRNA and protein (Figure 2A–C), JAM-A surface expression (Figure 2D), histological assessment of septal thickness (Figure 2E–I), total protein levels in BAL fluid (Figure 2J), and Evans blue extravasation (Figure 2K–L).

**Figure 2:**
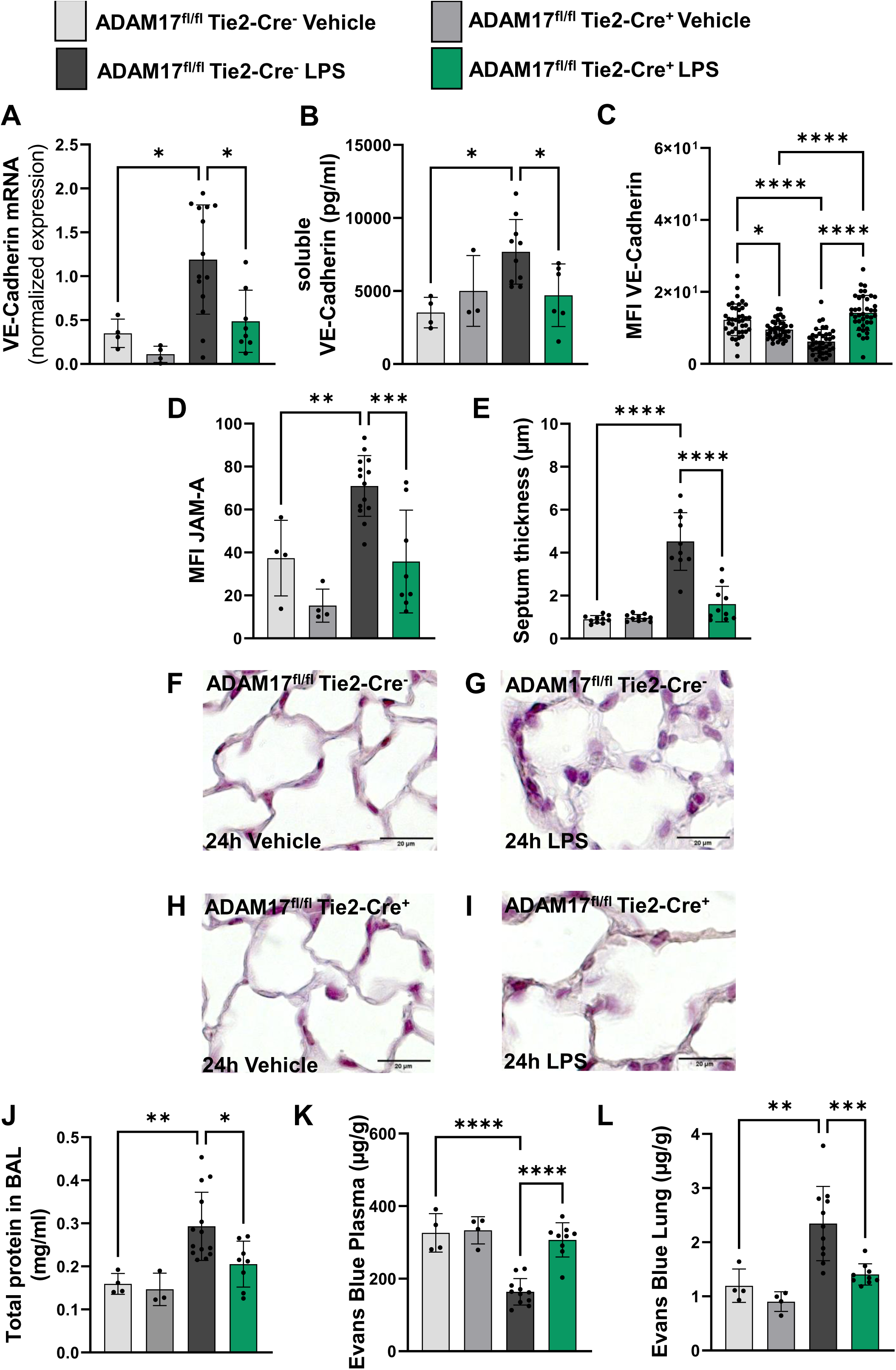
Endothelial ADAM17 promotes endothelial barrier disruption during lung injury. **(A)** Gene expression of vascular endothelial (VE)-cadherin in lung tissue three hours after LPS inhalation, assessed by RT-qPCR (n = 4/4/14/8 mice per group). **(B)** Quantification of soluble VE-cadherin in bronchoalveolar lavage (BAL) fluid three hours after LPS inhalation by ELISA (n = 4/3/10/6). (C) Quantification of VE-cadherin MFI in lung tissue using LasX (n = 4 mice per group; 10 measurements per image). **(D)** Quantification of junctional adhesion molecule-A (JAM-A) MFI on pulmonary endothelial cells 24 hours after LPS inhalation (n = 4/4/14/8). **(E)** Quantification of alveolar septal thickness using ImageJ (n = 4 mice per group; 10 measurements per group). **(F–I)** Representative histological images illustrating alveolar septal thickness under the indicated conditions (original magnification ×20). **(J)** Total protein concentration in BAL fluid 24 hours after LPS inhalation (n = 4/3/14/8). **(K–L)** Photometric quantification of Evans blue in plasma (K) and lung tissue (L) six hours after LPS inhalation (n = 4/4/11/9). Data are presented as mean ± SD. Statistical analysis was performed using one-way ANOVA with appropriate post hoc testing.

Three hours after LPS inhalation, ADAM17^fl/fl^ Tie2-Cre^−^ mice showed increased VE-Cadherin mRNA expression (Figure 2A) and elevated soluble VE-Cadherin (sVE-Cadherin) in BAL fluid (Figure 2B), indicating enhanced cleavage. In contrast, ADAM17^fl/fl^ Tie2-Cre^+^ knockout mice exhibited reduced transcription, diminished sVE-Cadherin release, preserved endothelial VE-Cadherin expression (Figure 2C), and fewer intercellular gaps (Supplementary Figure 3A–D), suggesting improved junctional integrity.

JAM-A surface expression on pulmonary endothelial cells was quantified by flow cytometry 24 hours after LPS exposure. LPS significantly increased JAM-A mean fluorescence intensity (MFI) in control mice, whereas endothelial ADAM17 deficiency markedly reduced JAM-A surface expression (Figure 2D).

Lung histology demonstrated substantial architectural preservation in knockout mice. Mayer’s hematoxylin staining revealed significantly reduced septal thickness compared to controls (Figure 2E), supported by representative images (Figure 2F–I). Total protein concentration in BAL fluid, measured 24 hours after LPS inhalation, was significantly lower in knockout mice compared to controls, indicating reduced alveolar-capillary leakage (Figure 2J).

Finally, we quantified microvascular leakage using the Evans blue extravasation assay six hours after LPS exposure, corresponding to the peak leakage time in murine acute lung injury (2). We observed reduced plasma Evans blue concentrations in control but not in knockout mice (Figure 2K), while knockout lungs showed significantly less Evans blue accumulation (Figure 2L), indicating preserved endothelial barrier function.

### Endothelial ADAM17 enhances neutrophil migration into the inflamed lung

Next, we investigated how endothelial ADAM17 affects neutrophil infiltration during LPS-induced lung inflammation. Representative immunohistochemical images (Figure 3A–D) show neutrophils, identified as brown-stained cells (red arrows), migrating into lung tissue. Quantification revealed a significantly lower number of neutrophils in the lungs of ADAM17^fl/fl^ Tie2-Cre^+^ knockout mice compared to ADAM17^fl/fl^ Tie2-Cre^−^ controls (Figure 3E).

**Figure 3:**
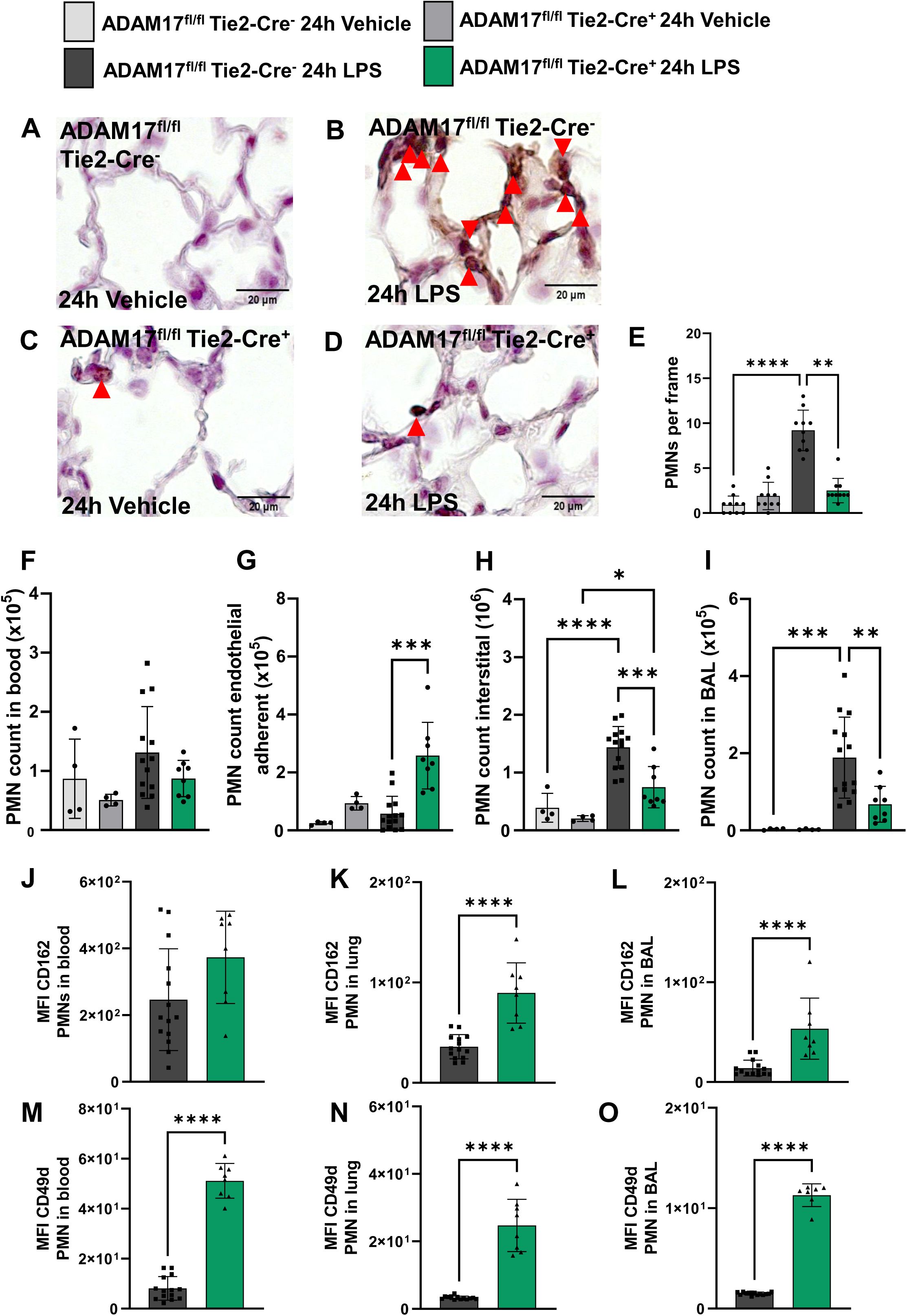
**Endothelial ADAM17 enhances neutrophil migration into the inflamed lung.**(A–D) Representative immunohistochemical images showing polymorphonuclear neutrophil (PMN) distribution in lung tissue under the indicated conditions. PMNs were stained using an anti-Ly6G+C antibody (brown) with hematoxylin counterstain. Red arrows indicate PMNs (n = 4 mice per group; original magnification ×20). (E) Quantification of PMNs per high-power field (HPF; n = 4 mice per group, 10 images per group). (F–I) Flow cytometric quantification of PMNs in blood (F), adherent to the pulmonary endothelium (G), within the interstitium (H), and in BAL fluid (I) 24 hours after LPS inhalation (n = 4/4/12–14/8). (J–L) Quantification of CD162 (PSGL-1) MFI and (M–O) CD49d MFI on PMNs isolated from blood, lung, and BAL Data are presented as mean ± SD. Statistical analysis was performed using one-way ANOVA.

To further dissect neutrophil migration across pulmonary compartments, we performed compartment-specific flow cytometry analysis 24 hours after LPS exposure, corresponding to peak transendothelial and transepithelial migration (2). We quantified neutrophils in blood (Figure 3F), adhered to the pulmonary endothelium (Figure 3G), within the interstitium (Figure 3H), and in the alveolar space (Figure 3I). Circulating neutrophil counts were similar between knockout and control mice (Figure 3F). However, knockout mice showed significantly more neutrophils retained at the vascular endothelium (Figure 3G) and significantly fewer neutrophils in the interstitium (Figure 3H) and alveolar space (Figure 3I), indicating impaired transmigration.

We next assessed whether endothelial ADAM17 affects neutrophil adhesion molecule expression. Flow cytometry analysis of PMNs in blood, lung, and BAL fluid revealed significantly elevated surface expression of CD162 (Figures 3J–L) and CD49d (Figures 3M–O) in knockout mice. CD162 was increased in lung and BAL PMNs, while CD49d expression was significantly upregulated in all compartments compared to controls. This upregulation of CD162 and CD49d correlates with the enhanced retention of PMNs at the endothelium and reduced tissue infiltration.

Together, these data demonstrate that endothelial ADAM17 promotes neutrophil transmigration into inflamed lung tissue by modulating adhesion molecule dynamics and vascular barrier interactions.

### Endothelial ADAM17 selectively amplifies cytokine receptor signaling

We next examined whether endothelial ADAM17 regulates inflammatory signaling pathways during LPS-induced lung inflammation. Using gene expression analysis, we assessed ADAM17 maturation and transport factors, TNFR and IL-6R signaling cascades, and TLR4-dependent pathways (Figure 4).

**Figure 4:**
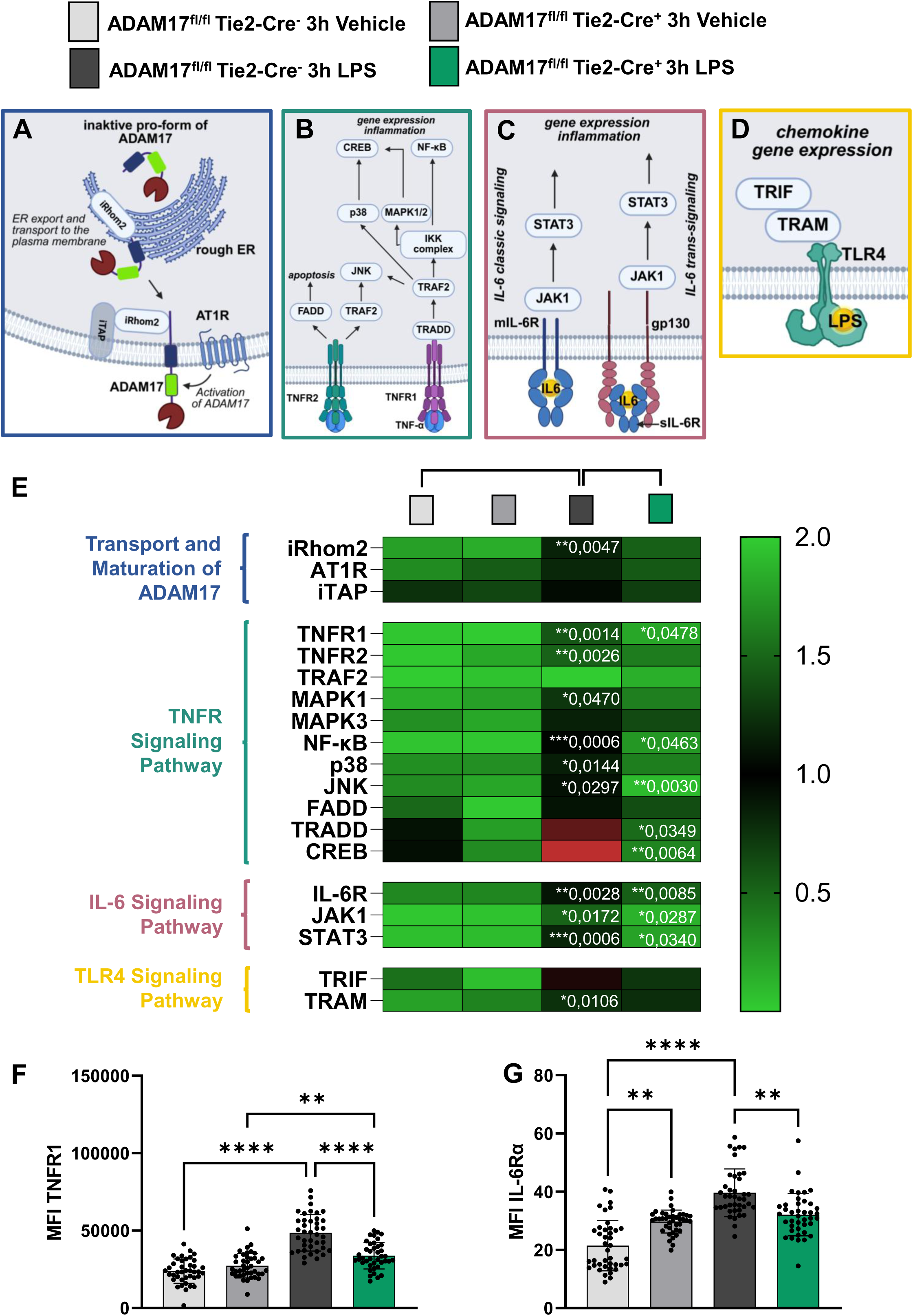
Endothelial ADAM17 amplifies inflammatory TNFR1 and IL-6R signaling pathways in acute lung injury. **(A)** Schematic overview of ADAM17 maturation and intracellular transport. **(B–D)** Schematic representations of TNFR1/2 **(B)**, IL-6R **(C)**, and TLR4 **(D)** signaling pathways. **(E)** RT-qPCR-based gene expression analysis of inflammatory signaling components in murine lung tissue three hours after LPS inhalation (n = 4/4/10–14/6–8). **(F–G)** Quantification of TNFR1 **(F)** and IL-6Rα **(G)** MFI in lung tissue using LasX (n = 4 mice per group; 10 measurements per image). Data are presented as mean ± SD; Statistical analysis: *P < 0.05, **P < 0.01, ***P < 0.001, ****P < 0.0001, using one-way ANOVA.

Three hours after LPS inhalation, expression of the ADAM17 regulatory proteins iRhom2, AT1R, and iTAP increased similarly in control and knockout mice, indicating that endothelial ADAM17 does not influence ADAM17 maturation or trafficking (Figure 4A).

In contrast, endothelial ADAM17 deletion selectively attenuated proinflammatory receptor signaling. Knockout mice showed a significant reduction in TNFR1 mRNA expression compared to controls, while TNFR2 expression remained unchanged (Figure 4B). Consistent with impaired TNFR1 signaling, transcript levels of downstream mediators Nf-κB, JNK, TRADD, and CREB were significantly reduced. Endothelial ADAM17 deletion also significantly decreased IL-6R expression, accompanied by reduced induction of the downstream signaling molecules JAK1 and STAT3 (Figure 4C).

By contrast, LPS activated the TLR4 pathway independently of endothelial ADAM17. Expression of the adaptor molecules TRIF and TRAM increased after LPS exposure to a similar extent in knockout and control mice, indicating preserved TLR4 signaling (Figure 4D).

We confirmed these transcriptional findings at the protein level. Immunofluorescence analysis 24 hours after LPS inhalation revealed significantly reduced expression of TNFR1 and IL-6Rα in lung tissue of knockout mice compared to controls (Supplementary Figure 4; Figure 4F–G). Notably, under basal conditions, knockout mice exhibited higher IL-6Rα expression than controls, suggesting altered receptor homeostasis in the absence of endothelial ADAM17.

Together, these data demonstrate that endothelial ADAM17 selectively amplifies TNFR1- and IL-6R-dependent inflammatory signaling while leaving ADAM17 maturation and TLR4 activation intact, positioning endothelial ADAM17 as a key modulator of cytokine-driven amplification loops in acute pulmonary inflammation.

### Endothelial ADAM17 amplifies alveolar inflammatory mediator release

We next examined how endothelial ADAM17 influences alveolar inflammation by quantifying cytokines and neutrophil-derived mediators in bronchoalveolar lavage (BAL) fluid following LPS exposure (Figure 5).

**Figure 5:**
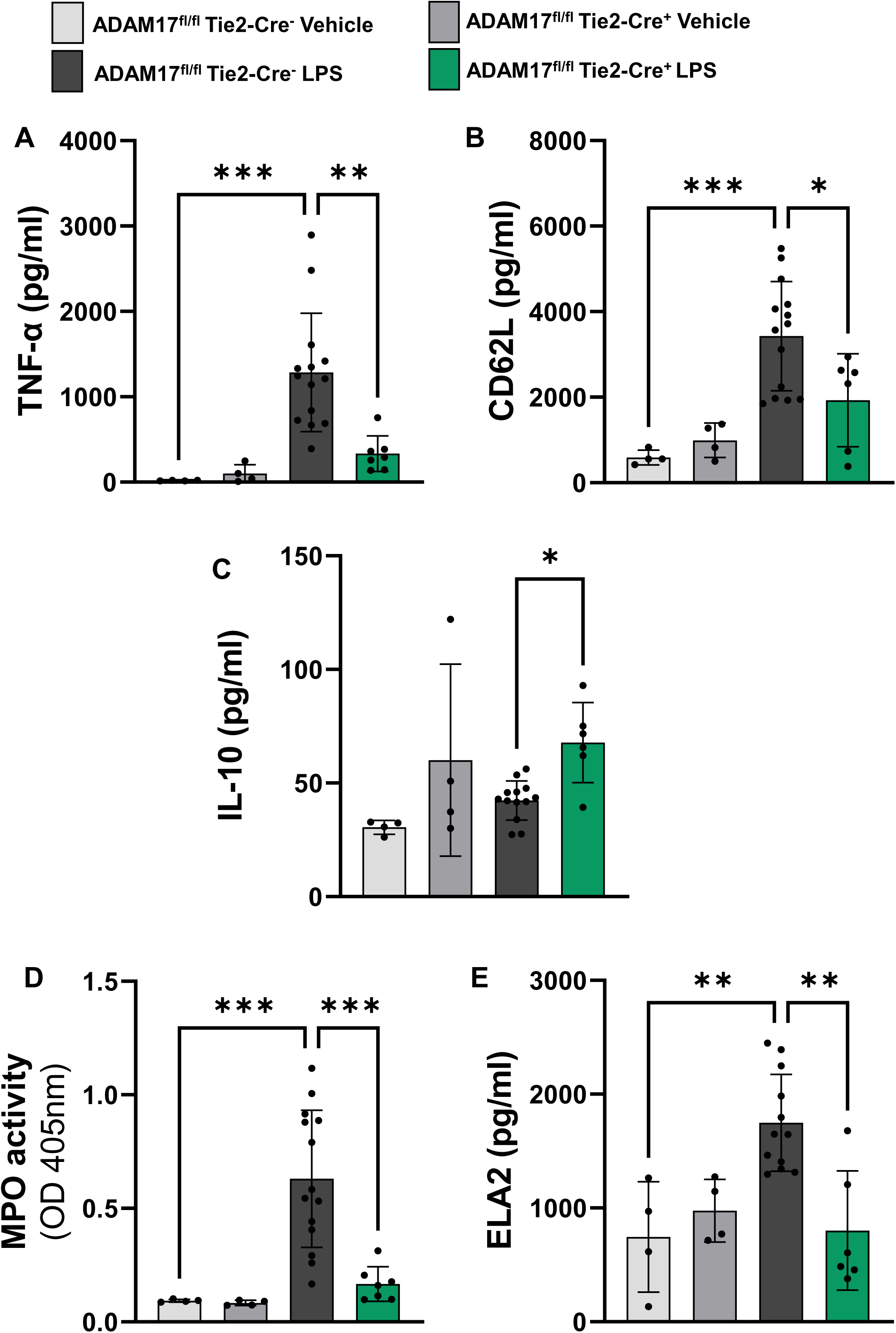
Endothelial ADAM17 increases alveolar inflammation during acute lung injury. Quantification of (A) TNF-α, (B) soluble L-selectin (CD62L), (C), Interleukin-10 (IL-10) three hours after LPS inhalation in bronchoalveolar lavage (BAL) by ELISA. Evaluation of neutrophil activity by (D) semiquantitative analysis of myeloperoxidase (MPO) and (E) quantification of neutrophil elastase (ELA2) by ELISA 24 hours after LPS exposure in BAL fluid. Data are presented as mean ± SD (n=4/4/12-14/6-7); Statistical analysis: *P < 0.05, **P < 0.01, ***P < 0.001, ****P < 0.0001, using one-way ANOVA.

Three hours after LPS inhalation, control mice exhibited a pronounced increase in TNF-α levels in BAL fluid, whereas endothelial ADAM17 deletion significantly reduced TNF-α release (Figure 5A). This reduction indicates impaired shedding of membrane-bound TNF-α and attenuated generation of soluble TNF-α in the absence of endothelial ADAM17.

LPS stimulation also markedly increased the release of soluble CD62L in control mice, while knockout mice showed significantly lower sCD62L levels in BAL fluid (Figure 5B). Reduced CD62L shedding coincided with enhanced retention of neutrophils at the vascular endothelium observed in vivo, consistent with impaired transmigration.

In contrast, knockout mice displayed significantly higher levels of the anti-inflammatory cytokine IL-10 following LPS exposure compared to controls, indicating a shift toward an anti-inflammatory alveolar milieu (Figure 5C).

Markers of neutrophil activation further supported this protective phenotype. Control mice showed robust increases in myeloperoxidase (MPO) and neutrophil elastase (ELA2) in BAL fluid 24 hours after LPS inhalation, whereas endothelial ADAM17 deletion markedly attenuated the release of both enzymes (Figure 5D–E).

Together, these findings demonstrate that endothelial ADAM17 promotes alveolar inflammation by enhancing the release of proinflammatory cytokines, facilitating CD62L shedding, and driving neutrophil activation, while its deletion limits alveolar tissue injury and favors an anti-inflammatory environment.

### Systemic ADAM17 inhibition attenuates LPS-induced lung inflammation

We next investigated whether pharmacological inhibition of ADAM17 recapitulates the protective phenotype observed in endothelial-specific knockout mice. We analyzed neutrophil migration across pulmonary compartments 24 hours after LPS inhalation following systemic inhibition with TAPI-1 or GW280264X (Figure 6).

**Figure 6:**
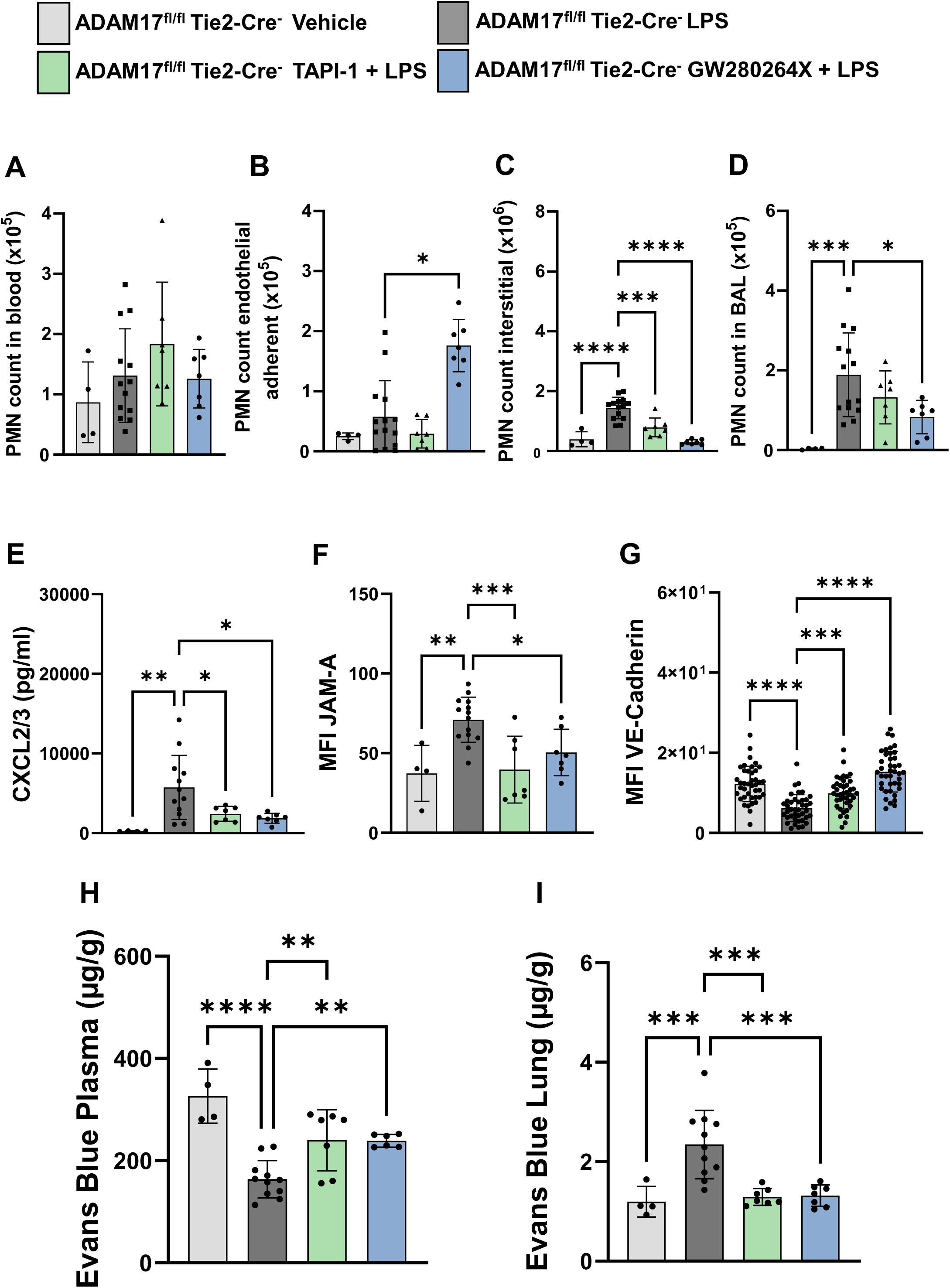
Systemic ADAM17 inhibition reduces LPS-induced lung inflammation. (A–D) Flow cytometric analysis of PMN distribution across lung compartments—blood **(A)**, endothelial-adherent **(B)**, interstitial **(C)**, and BAL **(D)** -24 hours after LPS inhalation with or without systemic ADAM17 inhibition (n = 4/13–14/7/7). **(E)** Quantification of CXCL2/3 in BAL fluid three hours after LPS inhalation by ELISA (n = 4/12/7/7). **(F)** Quantification of JAM-A MFI on pulmonary endothelial cells 24 hours after LPS inhalation (n = 4/14/7/7). **(G)** Quantification of VE-cadherin MFI in lung tissue using LasX (n = 4 mice per group; 10 measurements per image). **(H–I)** Photometric quantification of Evans blue in plasma **(H)** and lung tissue **(I)** six hours after LPS inhalation (n = 4/11/7/6–7). The data shown for ADAM17^fl/fl^ Tie2-Cre^-^ controls were previously presented in the following panels: Fig. 6A–D in Fig. 3F–I, Fig. 6F in Fig. 2D, Fig. 6G in Fig. 2C, and Fig. 6H–I in Fig. 2K–L (to minimize animal numbers in accordance with the 3R principle). Data are presented as mean ± SD; Statistical analysis: *P < 0.05, **P < 0.01, ***P < 0.001, ****P < 0.0001, using one-way ANOVA.

LPS exposure markedly increased neutrophil migration from the circulation into the lung interstitium and alveolar space (Figure 6A–D). TAPI-1 treatment significantly reduced neutrophil accumulation in the interstitium, while GW280264X treatment decreased neutrophil numbers in both the interstitium and alveoli. Combined inhibition of ADAM17 and ADAM10 further potentiated this effect, suggesting additive or synergistic roles of these proteases. Notably, dual inhibition led to a significant accumulation of neutrophils at the pulmonary endothelium, indicating impaired transendothelial migration and retention of immune cells at the vascular wall.

Systemic ADAM17 inhibition also attenuated inflammatory signaling and endothelial junction remodeling. To determine whether systemic ADAM17 inhibition additionally modulates neutrophil-attracting chemokine signals within the alveolar compartment, we quantified CXCL2/3 in BAL fluid. Pre-treatment with TAPI-1 or GW280264X significantly reduced LPS-induced release of the neutrophil-attracting chemokines CXCL2/3 (Figure 6E). LPS increased JAM-A surface expression on pulmonary endothelial cells, whereas inhibition of ADAM17 alone or in combination with ADAM10 significantly reduced JAM-A mean fluorescence intensity (Figure 6F). In parallel, LPS reduced VE-Cadherin expression and increased intercellular gap formation, while systemic inhibition of ADAM17 restored VE-Cadherin levels and preserved endothelial junction integrity (Figure 6G; Supplementary Figure 3).

Finally, we assessed microvascular permeability using the Evans blue extravasation assay six hours after LPS exposure. LPS significantly increased Evans blue accumulation in lung tissue, whereas systemic ADAM17 inhibition markedly reduced dye extravasation (Figure 6I). Consistent with reduced vascular leakage, plasma Evans blue concentrations increased following ADAM17 inhibition compared to LPS-treated controls (Figure 6H).

Together, these data demonstrate that systemic ADAM17 inhibition attenuates key pathophysiological hallmarks of ARDS, including excessive neutrophil recruitment, endothelial barrier disruption, and vascular leakage, thereby supporting ADAM17 as a rational therapeutic target in acute lung injury.

## Discussion

In this study, we identify endothelial ADAM17 as a central regulator of neutrophil recruitment, endothelial barrier integrity, and inflammatory signal amplification during LPS-induced acute pulmonary inflammation. By combining endothelial-specific genetic deletion with systemic pharmacological inhibition, we demonstrate that ADAM17 expressed by the pulmonary endothelium drives key pathophysiological hallmarks of ARDS, including excessive neutrophil transmigration, junctional destabilization, vascular leakage, and alveolar inflammation. Together, our findings position endothelial ADAM17 as a critical inflammatory amplifier and a rational therapeutic target in acute lung injury.

LPS exposure induced a pronounced upregulation of ADAM17 mRNA and protein expression in lung tissue. Although the transcriptional regulation of ADAM17 during acute pulmonary inflammation remains incompletely understood, increased expression likely represents an additional layer of inflammatory control beyond post-translational activation. Elevated ADAM17 levels can amplify inflammation by promoting the release of cytokines, cytokine receptors, and adhesion molecules. Consistent with our findings, ADAM17 expression is increased in chronic inflammatory lung diseases such as asthma and COPD (3), and LPS augments both ADAM17 transcription and shedding activity in endothelial cells (4). Notably, LPS selectively increased ADAM17 but not ADAM10 transcription, reinforcing the dominant role of ADAM17 in endotoxin-driven inflammation (4).

To specifically dissect the endothelial contribution, we employed the conditional ADAM17^fl/fl^ Tie2-Cre^+^ mouse model. Endothelial ADAM17 deletion reduced pulmonary ADAM17 mRNA expression by 77.5%, confirming that endothelial cells constitute a major source of ADAM17 in inflamed lung tissue. These findings are consistent with previous reports in Tie2-adam17^-/-^ mice (4). Although the Tie2 promoter has been debated for activity in hematopoietic cells (5), extensive characterization has demonstrated endothelial-restricted deletion of ADAM17 with preserved expression in peripheral blood cells and alveolar macrophages (4, 6). Our flow cytometric validation further confirmed intact ADAM17 expression in PMNs and platelets, supporting endothelial specificity.

Endothelial ADAM17 exerted profound effects on endothelial barrier regulation and microvascular permeability. In control mice, LPS caused severe architectural lung damage, septal thickening, and protein-rich pulmonary oedema, whereas endothelial ADAM17 deficiency preserved lung structure and barrier integrity. These findings mirror earlier observations by Dreymueller et al., who demonstrated protection from endotoxin-induced lung injury in Tie2-adam17^-/-^ mice (4).

At the molecular level, endothelial ADAM17 regulated key junctional proteins. JAM-A, a critical tight-junction component under physiological conditions (7), redistributes to the apical endothelial surface during inflammation to facilitate leukocyte transmigration (8, 9). This redistribution and increased JAM-A surface expression were absent in endothelial ADAM17-deficient mice. Previous studies demonstrated that LPS induces junctional disruption through shedding of JAM-A and cadherins (10), a process attenuated by ADAM17 inhibition (4, 11). Our data extend these findings by directly linking endothelial ADAM17 to JAM-A regulation *in vivo*.

VE-Cadherin represents a second critical junctional target. Although ADAM10 is considered the primary VE-Cadherin sheddase (11, 12), ADAM17 may contribute under specific inflammatory conditions (6). Reduced soluble VE-Cadherin release and preserved surface expression in endothelial ADAM17-deficient mice indicate that ADAM17 promotes junctional destabilization and gap formation during acute inflammation. Whether this reflects direct cleavage or secondary effects of dampened inflammation remains an important question for future studies.

Functional permeability assays supported these structural findings. LPS-induced increases in BAL protein concentration and Evans blue extravasation were abolished in endothelial ADAM17-deficient mice. While BAL protein content alone incompletely reflects barrier function (13), direct assessment of vascular permeability by Evans blue provides compelling evidence that endothelial ADAM17 critically governs microvascular leakage. To our knowledge, this represents the first direct demonstration of reduced Evans blue extravasation in ADAM17^fl/fl^ Tie2-Cre^+^ mice.

Endothelial ADAM17 also emerged as a key regulator of neutrophil trafficking. Endothelial ADAM17 deficiency caused pronounced retention of PMNs at the vascular endothelium with impaired transmigration into the interstitium and alveolar space, accompanied by increased expression of CD49d and PSGL-1. Dreymueller et al. similarly reported reduced pulmonary leukocyte recruitment in Tie2-adam17-/- mice (4), and histological analyses implicated shedding of adhesion molecules such as JAM-A and CX3CL1. ADAM17 cleaves multiple endothelial adhesion molecules, including ICAM-1, VCAM-1, JAM-A, and PECAM-1, thereby regulating adhesion– deadhesion dynamics (14). In contrast to endothelial deletion models, most studies on ADAM17 in leukocyte migration have focused on hematopoietic cells (15–18), highlighting the novelty of our endothelial-centric approach.

At the signaling level, endothelial ADAM17 selectively amplified TNFR1 and IL-6R pathways. Endothelial ADAM17 deletion reduced TNFR1 gene and protein expression and attenuated downstream signaling via Nf-κB, JNK, TRADD, and CREB. ADAM17 is well established as the principal sheddase for TNF-α and TNFR1 (19, 20), and recent work implicates ADAM17-dependent TNFR1 shedding in endothelial necroptosis and transendothelial migration (21). The reduced TNFR1 expression observed here likely reflects feedback inhibition secondary to diminished inflammatory signaling rather than direct transcriptional control.

Similarly, endothelial ADAM17 deletion attenuated IL-6R signaling, reducing IL-6R, JAK1, and STAT3 expression. ADAM17-mediated IL-6R shedding enables IL-6 trans-signaling (22–25), and diminished STAT3 activation in ADAM17-deficient settings has been reported previously (26). Compared with leukocyte-specific ADAM17 deletion, which only modestly reduces alveolar IL-6R levels (15), our data underscore the dominant role of endothelial ADAM17 in pulmonary IL-6 signaling.

Endothelial ADAM17 further amplified alveolar inflammation by promoting release of TNF-α, soluble CD62L, MPO, and ELA2. Although TNF-α production was reduced, it was not abolished, consistent with contributions from epithelial cells and alveolar macrophages (27) and alternative proteases (28). Notably, endothelial ADAM17 deletion reduced CD62L shedding, a process traditionally attributed to leukocyte-derived ADAM17 (29–31). Our findings suggest that endothelial ADAM17 contributes to CD62L shedding during leukocyte–endothelial interactions, revealing a previously underappreciated regulatory axis.

Reduced MPO and ELA2 release further indicated attenuated neutrophil activation. Although these enzymes are released by neutrophils independently of endothelial ADAM17 activity, their reduction likely reflects diminished neutrophil recruitment and activation within the alveolar compartment. This attenuation may limit excessive ROS production, NET formation, and tissue injury (32–34).

Importantly, systemic ADAM17 inhibition phenocopied the endothelial-specific knockout. Pharmacological inhibition reduced neutrophil recruitment, stabilized endothelial junctions, decreased chemokine release, and preserved vascular barrier integrity. Dual inhibition of ADAM17 and ADAM10 further enhanced these effects, suggesting cooperative protease activity. These findings align with prior studies demonstrating reduced PMN recruitment and permeability following ADAM17 inhibition (4, 21, 35) and extend them by integrating genetic and pharmacological evidence.

Collectively, our study establishes endothelial ADAM17 as a pivotal mediator of ARDS pathogenesis, orchestrating neutrophil trafficking, endothelial barrier disruption, and inflammatory amplification. By attenuating multiple interconnected pathophysiological processes, endothelial ADAM17 emerges as a promising therapeutic target for limiting excessive pulmonary inflammation and vascular injury in acute lung injury.

## Acknowledgments

Tie2-Cre mice (Tg(Tek-cre)12Flv; MGI ID: MGI: 2136412) were kindly provided by Prof. Dr. Peter Rosenberger (Department of Anesthesiology and Intensive Care Medicine, University Hospital of Tuebingen, Germany).

This work was supported by the German Research Foundation (Deutsche Forschungsgemeinschaft, DFG; grant KO 4280/4-1 to M.K.)

